# Deep Visual Proteomics reveals DNA replication stress as a hallmark of Signet Ring Cell Carcinoma

**DOI:** 10.1101/2024.08.07.606985

**Authors:** Sonja Kabatnik, Xiang Zheng, Georgios Pappas, Sophia Steigerwald, Matthew P Padula, Matthias Mann

## Abstract

Signet Ring Cell Carcinoma (SRCC) is a rare and highly malignant form of adenocarcinoma with increasing incidence and poor prognosis due to late diagnosis and limited treatment options. We employed Deep Visual Proteomics (DVP), which combines AI directed cell segmentation and classification with laser microdissection and ultra-high sensitivity mass spectrometry, for cell-type specific proteomic analysis of SRCC across the bladder, prostate, liver, and lymph nodes of a single patient. DVP identified significant alterations in DNA damage response (DDR) proteins, particularly within the ATR and mismatch repair (MMR) pathways, indicating replication stress as a crucial factor in SRCC mutagenicity. Additionally, we observed substantial enrichment of immune-related proteins, reflecting high levels of cytotoxic T lymphocyte infiltration and elevated PD-1 expression. These findings suggest that pembrolizumab immunotherapy may be more effective than conventional chemotherapy for this patient. Our results provide novel insights into the proteomic landscape of SRCC, identifying potential targets and open up for personalized therapeutic strategies in managing SRCC.

## Introduction

Signet Ring (SR) cell carcinoma (SRCC) is a rare and highly aggressive type of adenocarcinoma that can occur in multiple organs. While the stomach is the most common primary tumor site, SRCC has also been reported in the prostate, breast, lung, and bladder ^1^. Regardless of origin it typically metastasizes rapidly to distal sites ^2,3^. Incidences of gastric SRCC have persistently increased over the last few decades ^4,5^.

If SRCC occurs from cells other than stomach glandular cells this may make disease classification in the effected organ more difficult ^6–8^. However, there is one pathological feature that characterizes SR cells as such: a high concentration of intercellular mucin that builds up in large vacuoles, pushing the nucleus to the periphery of the cell and giving it the distinctive shape of a signet ring ^9^.

Despite clinical advances in gastric cancer classification, grading and treatment, the SR cell carcinoma subtype remains a substantial clinical burden ^9,10^. Due to its rarity and a propensity for late symptom onset, SRCC patients are often diagnosed at an advanced stage, limiting treatment options and therapeutic efficacy ^11,12^. Surgical resection followed by postoperative chemotherapy and radiotherapy are the main management options for advanced disease^13^. However, these treatments have limited impact on overall survival and can have numerous negative effects that worsen patient wellbeing ^12^. The rarity of SRCC and the substantial knowledge gap regarding its fundamental biology and underlying signaling pathways thus combine to limit personalized therapeutic strategies for this distinct cancer subtype.

Investigations into SRCC biology have primarily revolved around this cancer’s inherently increased proliferation rate, characterized by aberrations of the RAS/RAF/MAPK^14^, HER2 or Wnt/β-catenin ^15^ signaling pathways and mutation of the E-cadherin gene CDH1 ^16^. Microsatellite instability and strong lymphocyte infiltration have also been linked with colorectal SRCC, clinicopathological signatures typically rather associated with colorectal cancer than specifically with SRCC ^17,18^. It is also kown that in colorectal SRCC, the SMAD complex triggers the epithelial-mesenchymal transition (EMT) in response to transforming growth factor (TGF)-β signaling, which accounts for the distinctive change of epithelial cell junctions and polarity in SRCC of the colon^19,20^.

So far, most research on SRCC has been limited to clinical observations, histological classifications ^21,22^, and obtaining genomic sequencing data specific to occurrences in affected organs ^20,23^, predominantly the colon. We reasoned that global molecular analyses at the protein level could contribute to elucidating the broader biological context and distinctive pathogenic mechanisms of SRCC. The spatial proteomics field has made significant strides in recent years, and is potentially able to address the above challenge ^24,25^. In particular our group has developed the Deep Visual Proteomics (DVP) technology which combines high-resolution image acquisition with machine learning-guided segmentation and classification, followed by single-cell type enriched high-sensitivity mass spectrometry (MS)-based proteomics^26^.

In this study, we took a precision oncology approach by using DVP to examine SRCC in four different organs—the bladder, prostate, liver, and lymph node—within a single patient. We reasoned that this spatial context would allow us to explore proteome differences and similarities of SR cells across tissues, offering valuable insights into tumor origin, potential mechanisms of metastasis and to make treatement recommendations.

## Results

### Patient Disease Background and Interventions

The patient was diagnosed with SRCC, with the bladder identified as the primary site of origin, following the removal of a suspicious mass on the bladder wall that was revealed by magnetic resonance imaging (MRI). Hematoxylin and eosin (H&E) staining of the mass revealed the typical signet ring morphology, and the patient was subjected to a radical cystectomy that removed the bladder (B.), prostate (P.), seminal vesicles (S.V.) and 14 lymph nodes (L.N.) (Figure 1A, B). Post-surgery pathology of these organs showed cells with signet ring morphology in all organs and nine out of fourteen lymph nodes. To enhance the therapeutic options for the patient, a genomic analysis was performed and a molecular tumor board report was filed, noting a microsatellite instability of only 0.8%, an ATRX (alpha thalassemia/mental retardation syndrome X-linked) frameshift mutation, MYCL and RICTOR (Rapamycin-insensitive companion of mTOR) amplification and KDM6A (Lysine-specific demethylase 6A) biallelic loss. The patient underwent chemotherapy with a combination of oxaliplatin, which was discontinued after four months due to the onset of continuous neuropathy, and capecitabine, likewise discontinued after seven months, before being monitored by quarterly computed tomography (CT) scans (Figure 1C). Twelve months after the cessation of chemotherapy, CT scan revealed suspicious enlargement of several lower abdominal lymph nodes. After further evaluation through a positron emission tomography (PET) scan, an accessible lymph node was removed by ultrasound guided biopsy, in which pathology confirmed the presence of cells of signet ring morphology. The patient received a combination of immunotherapy with pembrolizumab, which is ongoing, and chemotherapy with carboplatin, which was again stopped after four months due to side effects (Figure 1C). The tissues used in this study were obtained prior to any treatment.

**Figure 1.**
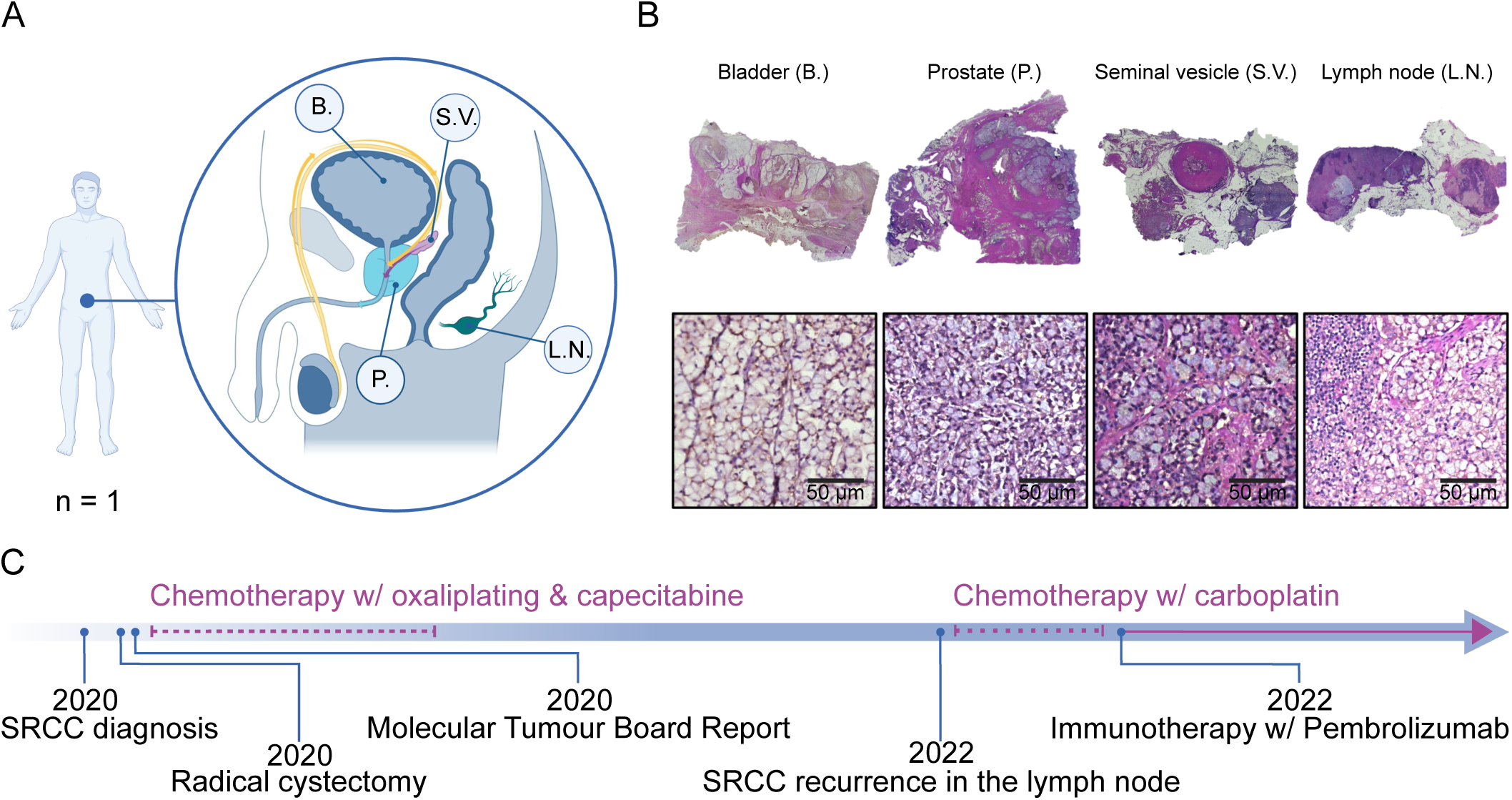
Signet ring cell carcinoma samples and timeline of medical interventions. **A** Sample overview of signet ring cell carcinoma (SRCC)-positive tissues including the bladder (B.), the seminal vesicle (S.V.), one lymph node (L.N.) and the prostate (P.). Image was adapted from <u>tulsaprocedure.com</u> and modified. **B** Images of hematoxylin and eosin (H&E) stained SRCC formalin-fixed, paraffin-embedded (FFPE) tissues. **C** Chronological timeline of medical interventions. Illustrated with <u>BioRender</u>.

### A simple stain allows robust segmentation and classification for the DVP workflow

For spatial proteomics we sectioned formalin-fixed, paraffin-embedded (FFPE) tissue blocks of all four organs (bladder, prostate, seminal vesicle and one lymph node) at three µm thickness using a microtome and mounted the tissue sections on polyethylene naphthalate (PEN) membrane-coated microscopy glass slides (Figure 2A). Tissues were stained with DAPI for nuclear visualization. A crucial step in DVP is delination of the cell plasma membrand for subsequent laser microdissection. For our samples, we found that staining by wheat germ agglutinin (WGA), a lectin that binds to specific carbohydrates in the plasma membrane, was sufficient for this purpose (Figure 2B). In comparison to other staining methods, such as cytokeratin 1 (CK1) or the conventional H&E, WGA staining proved superior in terms of efficiency and simplicity. Continuing with the DVP pipeline, we imaged the tissue slides with a standard immunoflourecent microscope (Zeiss Axio) and processed images with the Biology Image Analysis Software (BIAS) ^26^ (Figure 2A, B). For cell segmentation we fine-tuned a pre-trained model in BIAS. We trained a machine learning model for cell classification, which involved manual annotation of more than 1000 SRCC and lymphocytes from each organ to capture morphological diversity, ensuring accurate classification across tissue types (Figure 2B). Prediction accuracy of SR cells was 95% based on 10-fold cross validation and indendent validation by a pathologist.

**Figure 2.**
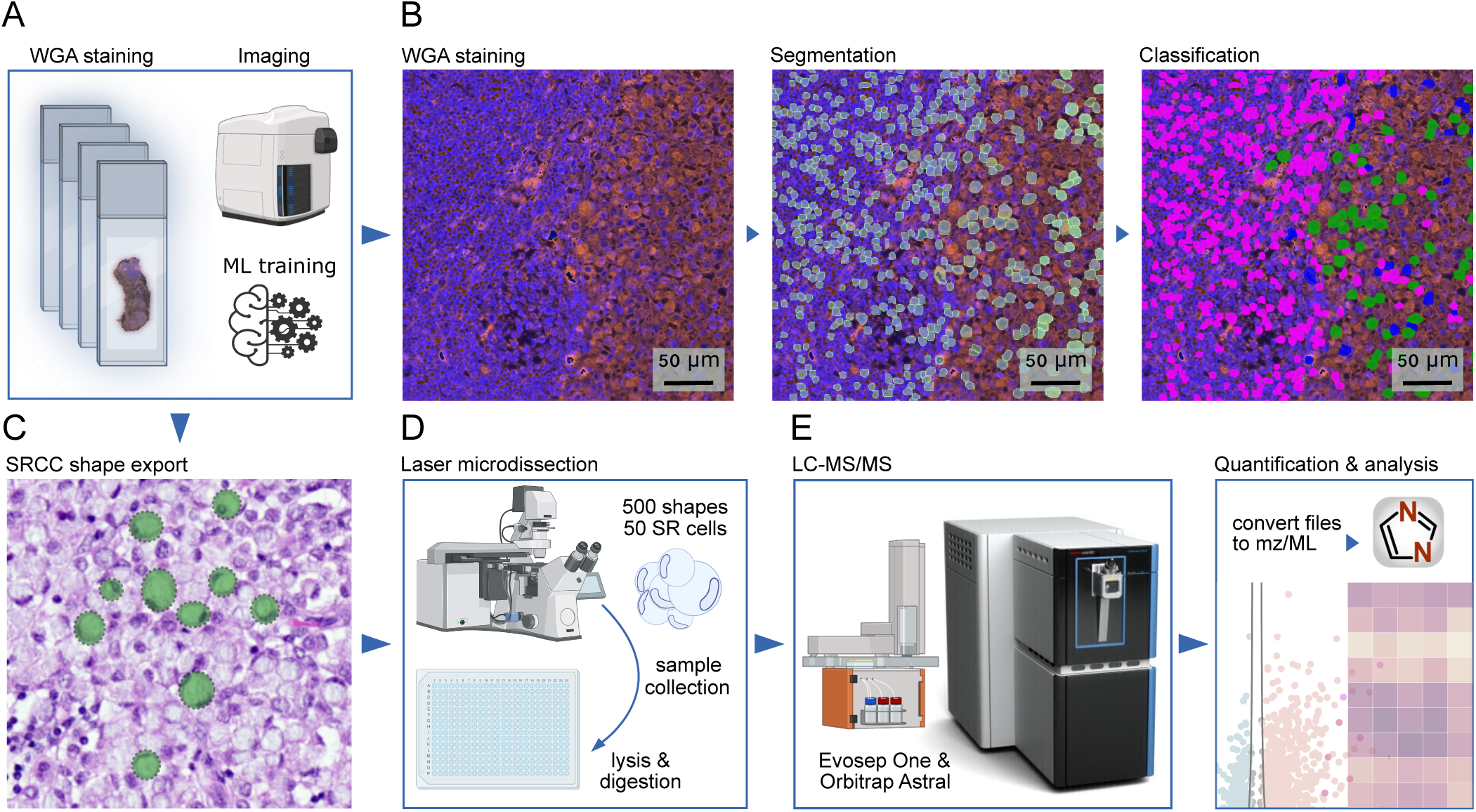
Deep Visual Proteomics workflow on WGA-stained tissues. **A** Cell-type specific tissue preparation for the Deep Visual Proteomics (DVP) spatial proteomics pipeline, starting with FFPE tissue sectioning, mounting, staining and image acquisition. **B** Representative images of WGA-stained lymph node tissue, showing one raw, one segmented and one classified image (lymphocytes in pink, SR cells in green and segmentation artifacts in blue). **C** Export mask of classified SR cells. **D** Illustration of the semi-automated laser microdissection sample collection and processing, followed by **E** the liquid chromatography-mass spectrometry (LC-MS) setup. Illustrated with <u>BioRender</u>.

Shapes were subsequently exported to a second microscope for semi-automated laser microdissection (Leica LMD7). In total, we dissected 500 cell shapes per organ, corresponding to approximately 50 SR cells, in triplicates. Collected cell shapes were lysed and enzymatically digested for subsequent MS-based proteomics (Figure 2D). Peptides were separated by to the Evosep One chromatography system ^27^, coupled to the Orbitrap Astral™ mass spectrometer ^24^. This was followed by protein identification and quantification using the DIA-NN software ^28^ (Figure 2E, see Methods).

### Proteomic analysis identifies organ-specific SRCC and DDR protein signatures

Analyzing MS data from the equivalent of 50 SR cells in all four organs, and including non-cancerous epithelial prostate cells as controls, we quantified a median of 6,638 different proteins (Figure 3A), with a excellent coefficient of variation (CV) of approximately 11% across the tissues (Figure 3C). A total of 4,648 proteins were present across all triplicates and organs and 7,157 in at least 70% of samples of each organ indicating high completeness of our data set (Figure 3C). Across the four organs, we identified 4,825 proteins as a common core proteome (Figure 3E). As expected, proteins uniquely present in each organ mirror specific organ functions, such as semenogelin-2 (SEMG2) in the seminal vesicle which is responsible for gel matrix formation for spermatozoa^29^ (Figure 3F).

**Figure 3.**
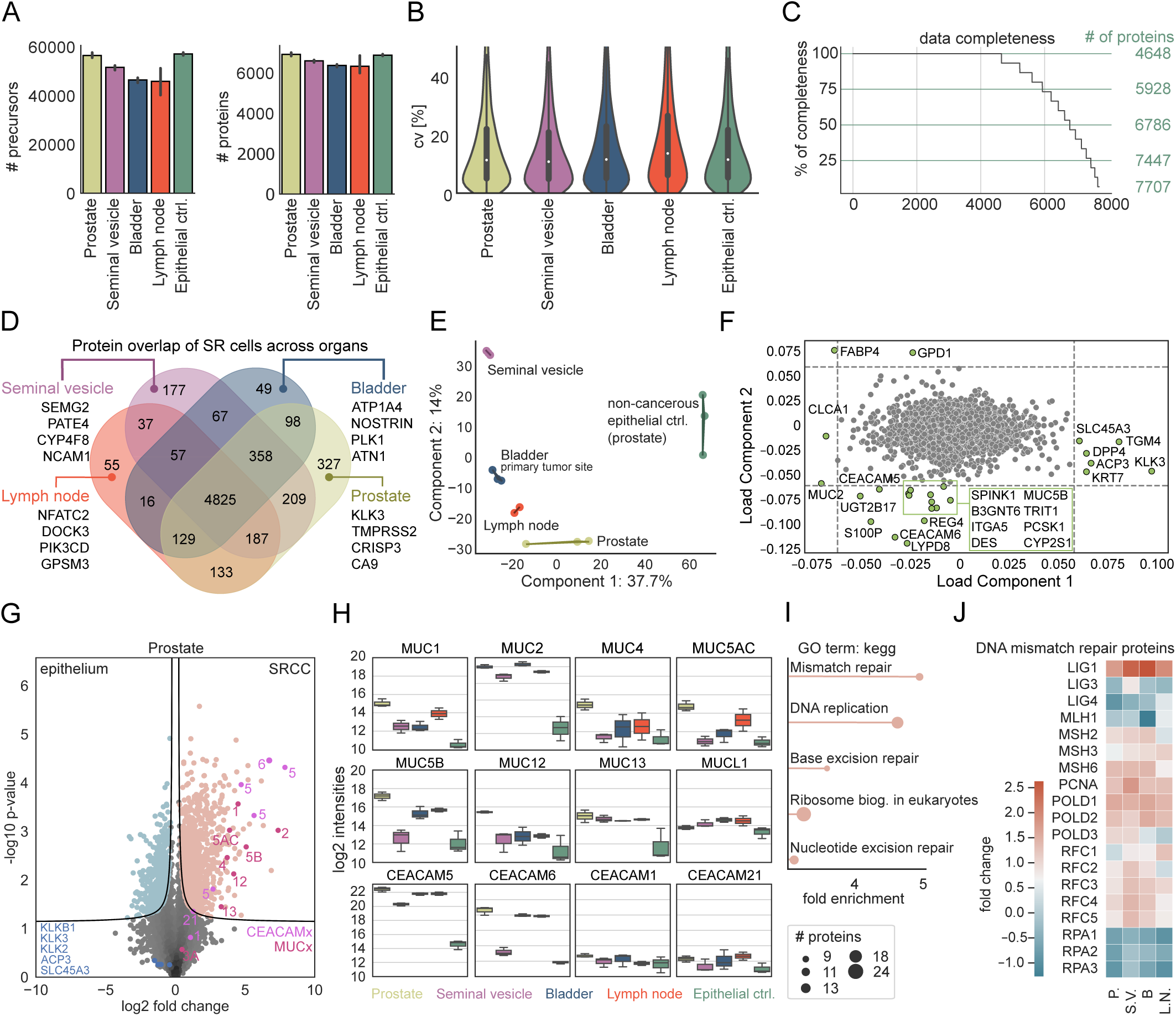
Proteomic depth and signatures of signet ring cells across tissues. **A** Number of precursors and proteins across all tissues. **B** Coefficients of variation (cv). **C** Data completeness and highlighted cutoffs at 100, 75, 50 and 25%. Proteins were ranked in descending order based on the number of valid values present across organs and triplicates. **D** Overlap of SR cell proteomes across tissues, highlighting organ-specific proteins for the seminal vesicle, bladder, lymph node, and prostate. **E** Principal component analysis (PCA) of SR proteins across tissues. **F** Loading plot of the PCA, highlighting outlier proteins. **G** Pairwise proteomic comparison of the non-cancerous epithelial control (Prostate ctrl.) cells to the SR cells of the prostate (two-sided t-test, FDR <0.01, s_0_ = 0.1). **H** Log2 normalized protein intensities of the mucin (MUC) and the carcinoembryonic antigen-related cell adhesion molecule (CEACAM) family members. **I** Gene Ontology (GO) term enrichment analysis using KEGG pathways of proteins significantly upregulated in the previous pairwise proteomic comparison. **J** Heatmap showing fold changes in DNA mismatch repair proteins between SR cells of the prostate, seminal vesicle, bladder, and lymph node to epithelial prostate cells as control.

Principle component analysis (PCA) clearly clustered samples originating from the same tissue, but also the cancerous SR cell away from the control (Figure 3E). Likewise, SR cells from the lymph node, prostate, and bladder were clearly distinct from SR cells of seminal vesicles (Figure 3E). Well known markers for prostate cancer and proteins involved in EMT including dipeptidyl peptidase 4 (DPP4), transglutaminase 4 (TGM4), keratin 7 (KRT7), acid phosphatase 3 (ACP3), kallikrein-related peptidase 3 (KLK3) and solute carrier family 45 member 4 (SLC45A4) ^30–34^ were among the proteins driving the separation between SRCC and epithelial control in our PCA along the load component 1 (Figure 3F). Proteins that are instead enriched in the SR cells compared to the epethilial control cells include carcinoembryonic antigen-related cell adhesion molecule 5 (CEACAM5) and CEACAM6, mucins (MUC2, MUC5B), and calcium-activated cloride channel regulator 1 (CLCA1), classical markers for SRCC (Figure 3F). Fatty acid binding protein 4 (FABP4) and glycerol-3-phosphate dehydrogenase 1 (GPD1) separate the SR cells from the seminal vesicle from the other organs through component 2, likely due to tissue-specific differences in cellular proteomes, function, due to interactions between SR cells and their tumor environment (Figure 3F). Thus, DVP recapitulated expected or recently described physiological patterns while adding novel molecular players.

In the prostate, there was a clear and significant enrichment of MUC and CEACAM proteins between the epithelial control and SR cells (Figure 3G). In contrast, we observed minimal differences in the levels of prostate and prostate cancer-associated proteins, including KLKB1, KLK2, KLK3, APC3, and SLC45A4, conventional adenocarcinoma of the prostate (Figure 3H). To control for SRCC-specific protein patterns and to investigate proteins with the most significant differential changes, we focused on two well-known protein families strongly associated with SRCC, mucins and CEACAMs.

MUC1, MUC2, and MUC13 showed the strongest – up to ten-fold - and most consistent enrichments in SR cells across all organs compared to the epithelial control cells of the prostate (Figure 3H). MUC1 and MUC2 are already well known to be overexpressed in gastric cancers, however, MUC13, a transmembrane mucin might play an yet unknown role in cell signaling and eptithelial barrier protection. MUC4, MUC5AC, MUC5B, and MUC12 had significant but fluctuating fold-changes between organs. SR cells in the seminal vesicles exhibited protein levels similar to those of non-cancerous control cells in the prostate. Mucin-like 1 protein (MUCL)1 has structural similarities and glycosylation patterns to classical mucins, but interestingly its expression profile was not significantly changed across all tissues analyzed, demonstrating that changes and overexpression in SR cells are specific to classical mucins.

Regarding the CEACAM family, CEACAM5 and CEACAM6 expression increased up to ten-fold between cancerous and epithelial controls, with the sole exception of CEACAM6 in the SR cells of the seminal vesicle. CEACAM1 and CEACAM21, who have different functions and structures, remained uniform across the different tissues supporting the notion that they are not directly involved in ^35^.

We next asked if the proteins highly enriched in prostate SR cells could point us to any therapeutically relevant pathways. Indeed, the top ones in terms of fold-change and statistical significance in a Gene Ontology (GO) enrichment analysis were all related to DNA replication and DNA damage response (DDR), including ‘nucleotide excision repair (NER)’, ‘base excision repair (BER)’ and ‘mismatch repair (MMR)’ (Figure 3I). We additionally found that the enrichment of MMR pathways is universal to all SR-positive tissues. The majority of the constituent proteins were upregulated, however, a number of prominent replication proteins (RPAs) were substantially downregulated (Figure 3J).

### Signet ring cells exhibit multiple DDR pathway deficiencies across organs

Following up on our observation that proteins of the DDR showed abundance changes between prostate SR cells and epithelial cells, we next investigated tissue-specific protein changes by correlating the fold-changes between them (Figure 4A). Comparing two tissues at a time, observed that Ly6/PLAUR domain-containing protein 8 (LYPD8) and UDP-glucuronosyltransferase 2B17 (UGT2B17) showed similar patterns to the above mentioned CEACAM5 and CEACAM6 proteins. LYPD8 is also involved in epethilial cell junction integrity, pointing to a dysregulation of cell-cell adhesion, as well as potential deficiencies in tissue protection. UGT2B17 is involved in the metabolism of steroid hormones and xenobiotics, which can alter the tumor microenvironment.

**Figure 4.**
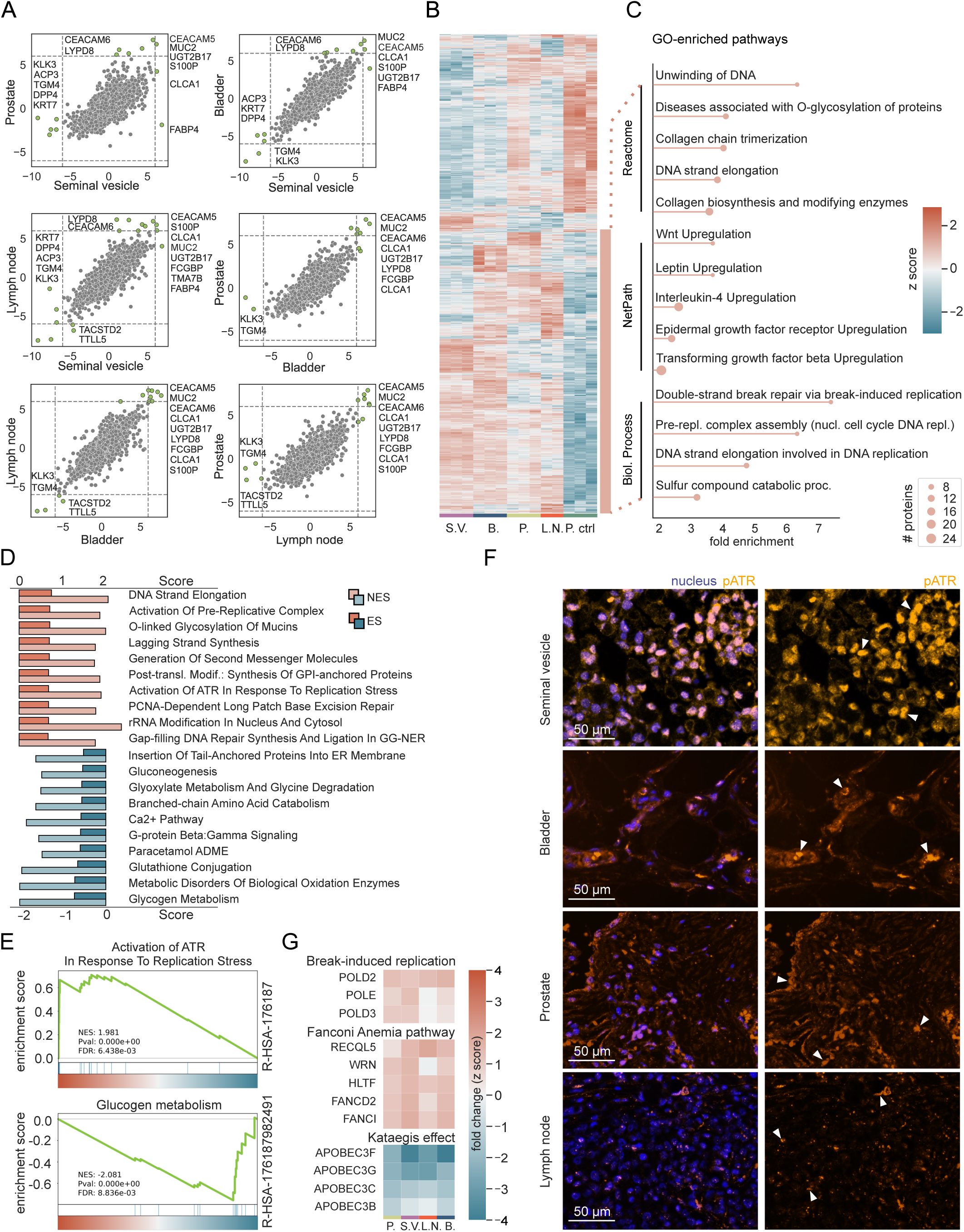
Proteomic profiling of signet ring cells in the context of DNA damage. **A** Inter-organ fold change correlation plots, with emphasis on significant protein variations highlighted in green (cutoffs at ±6 fold-change). **B** Unsupervised hierarchical clustering of ANOVA significant proteins (premutation-based FDR <0.01, s_0_ = 0.1). **C** GO term enrichment analysis of the bottom, upregulated cluster (in orange), highlighting the top five enriched pathways within Reactome, NetPath, and Biological Process. **D** Gene Set Enrichment Analysis (GSEA) of significantly positively and negatively enriched proteins after a pairwise proteomic comparison of SR cells to the epithelial cells of the prostate (two-sided t-test, FDR <0.01, s_0_ = 0.1). Top ten pathways, sorted in a descending sequence according to their enrichment score (ES), with the corresponding normalized enrichment score (NES). **E** Two representative GSEA graphs, showing one positively and one negatively enriched pathway. **F** Representative images of SRCC-positive regions of the seminal vesicle, bladder, prostate and lymph node, stained for pATR and DAPI (nucleus). The auto-fluorescence signal of the mucus was initially used to identify SRCC-positive tumor regions. The scale bar is at 50 µm, and white arrows indicate strong pATR accumulation with the nuclei. **G** Heatmaps showing fold changes of proteins involved in ‘break-induced replication’, the ‘Fanconi Anemia pathway’ and the ‘Kataegis effect’. SR cells of all four tissues (prostate, bladder, lymph node and the seminal vesicle) were compared to the epithelial cells of the prostate.

S100 calcium binding protein P (S100P), MUC2, and CLCA1 also had similar expression patterns across the tissue (Figure 4A), in line with CLCA1 (Calcium-activated chloride channel regulator 1) affecting mucin secretion through Ca^2+^ signalling and its possible implications in cancer pathophysiology^36^. KLK3 and TGM4, well-known prostate-specific markers, consistently exhibit a negative or zero fold change between tissues and non-cancerous control cells (Figure 4A). Thus signet ring cells may arise due to different molecular mechanisms distinct from those of conventional prostate adenocarcinoma and metastases.

To globally examine protein patterns prevalent across all SR cells and contrast them with epithelial cells as a control, we performed unsupervised hierarchical clustering on the 1,560 ANOVA significant proteins, which revealed two prominent clusters, those upregulated or downregulated with respect to control (upper, red cluster and lower, blue cluster in Figure 4B). We performed GO term enrichment analysis on the upregulated cluster using Reactome, NetPath, and Biological Processes, which highlighted diverse pathways active in SRCC cells. These included Wnt, leptin, epidermal growth factor (EGF) receptor and transforming growth factor β (TGFβ) receptor pathways, all well-known for their roles in various carcinomas including stomach, colorectal and SRCC, (Figure 4C). Apart from these, the most prominent pathways were again associated with DNA replication and DDR (Figure 4C).

Next, by comparing the SR cells to the epithelial control cells, we ran a gene set enrichment analysis (GSEA) on proteins which showed a significant enrichment following pairwise proteomic comparison. Remarkably, 7 of the top 10 pathways are part of DDR, namely ‘Activation of [the] pre-replicative complex’, ‘activation of ATR (ataxia-telangiectasia mutated and Rad3-Related), a pathway triggered upon perturbations affecting DNA replication dynamics characterized as replication stress (RS)’, ‘PCNA-dependent long patch base excision repair (LP-BER)’, ‘gap-filling DNA repair synthesis and ligation in global-genome nucleotide excision repair (GG-NER)’, and ‘DNA strand elongation’ (Figure 4D, E).

To validate our proteomic results regarding ATR signaling activation, we stained all SRCC-positive tissues for phospho-ATR (pATR), the activated form of the protein kinase which phosphorylates downstream key proteins involved in DDR ^37–40^. Our staining results confirmed the presence of pATR across our tissue samples, with the highest positivity observed in the seminal vesicle tissue (Figure 4F). We confirmed that the seminal vesicle is particularly highly positive for pATR whereas the bladder, prostate, and lymph node also display pATR signals, but to a lesser extent (Figure 4F).

Pathways implicated in metabolic processes such as ‘glycogen metabolism; and signaling mechanisms such as the ‘Ca^2+^ pathway’ and G-protein beta:gamma signaling’ are negatively enriched (Figure 4D, E). Downregulation of these pathways in SR cells likely reflects metabolic reprogramming of cancer cells, alterations in calcium signaling to support uncontrolled growth and survival, and specific adaptations of SRCC to facilitate mucin production and secretion. Given the observations of significant changes in protein abundances related to DDR pathways and the ATR signaling axis in SR cell-positive tissues compared to epithelial control cells, we further investigated proteins involved in stalled replication fork (RF) protection and repair of complex DNA lesions formed in case of replication fork collapse, a key part of the cellular response to DDR. These included proteins of the Fanconi Anemia (FA) pathway specifically the FA group D2 protein (FANCD2) and its interactor ^41^, Fanconi Anemia complementation group I (FANCI) ^42^. Additional mediators of the same process including DNA unwinding RecQ like helicase 5 (RECQL5), Werner syndrome helicase (WRN), and helicase-like transcription factor (HLTF) all displaying a similar positive fold change (Figure 4G). Our data provides strong indications of an ongoing RS and of the subsequent response of the SR cells to maintain their genomic stability by upregulating various RF protection mechanisms.

Upon persistent RS and prolonged RF stalling, replisome structure is impaired and RFs collapse, leading to the emergence of single-end double strand breaks (seDSBs), the most deleterious form of DNA lesions. Cells then trigger the highly error-prone break induced replication pathway (BIR) to deal with this threat ^43,44^. GSEA on our proteomic data showed a significant enrichment of this mutagenic pathway (Figure 4F). Moreover, DNA polymerase delta subunit POLD3, an essential subunit of DNA polymerase delta upon BIR, together with POLD2 and DNA polymerase epsilon (POLE) show a positive fold change enrichment comparing SR cells of the prostate, the seminal vesicle, the lymph node, and the bladder to the epithelial control (Figure 4G).

APOBEC3s, members of the Apolipoprotein B mRNA-editing enzyme catalytic polypeptides (APOBECs) superfamily, exhibit overexpression across various cancer types, notably bladder ^45–47^ and prostate cancer ^48,49^. The induced hyper-mutations of long stretches of single strand DNA (ssDNA) formed during BIR (with APOBEC3A and APOBEC3B being the major mutators) through deamination, foster genome instability in cancer cells, a phenomenon referred to as “kataegis”. However, proteins of the APOBEC3 family of enzymes were markedly reduced in abundance in SR cells of every tissue (Figure 4G), possibly as a protective feedback mechanism to mitigate the mutational burden and maintain genomic stability ^50–53^.

Collectively our analysis of the proteome changes of SR cells from the bladder, identified as the primary tumor site, as well as from metastatic sites, namely the prostate, seminal vesicle, and lymph node revealed consistent patterns of a severe dysregulation of multiple DNA repair mechanisms, with a potential negative impact on genome integrity.

### Enrichment of Complement System and PD-1 Signaling Proteins in Signet Ring Cells result in a cytotoxic T lymphocyte infiltration

DDR genes’ mutations and expression profiles have been recently associated with alterations of immune regulatory gene expression and CD8+ T cell infiltration in the tumor microenvironment, serving as a predictive marker of immune checkpoint blockade (ICB) therapy efficiency ^54,55^. We therefore hypothesized that our unique protein signatures could indicate a higher immunogenicity and a greater mutational burden of the SRCC. The Reactome-curated ‘Complement system’ pathway displayed a positive fold change across all tissues, with the most marked increases seen in the C1q subcomponent subunits A (C1QA), B (C1QB), and C (C1QC) (Figure 5A). A similar expression pattern was observed in immunoglobulins and proteins involved in the programmed cell death protein 1 (PD-1) signaling pathway (Figure 5A).

**Figure 5.**
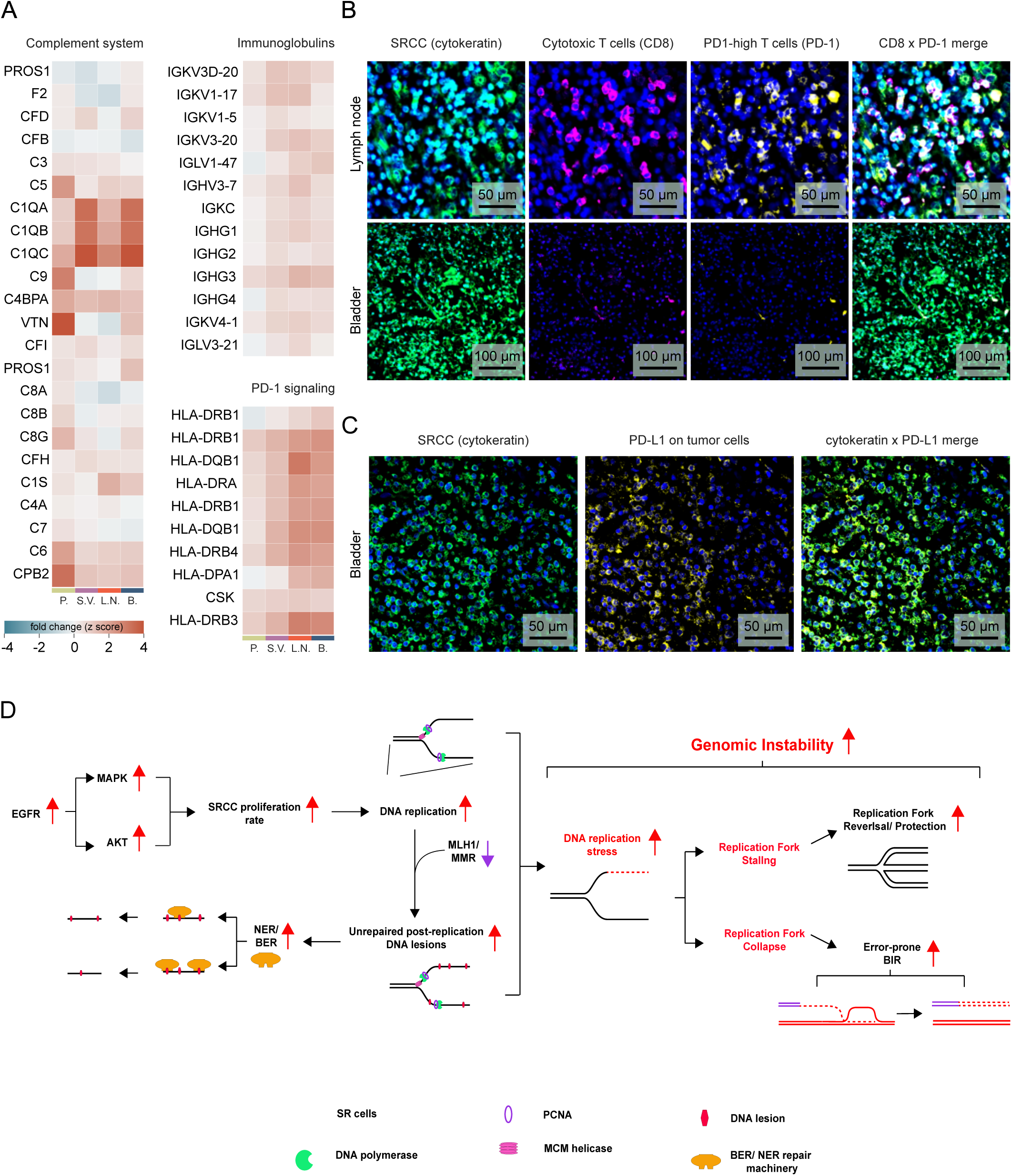
SRCC shows immunogenicity and cytotoxic T cell infiltration. **A** Fold-changes of proteins involved in GO ‘Complement system’ pathway, ‘immunoglobulins’ and ‘PD-1 signaling’. SR cells of all four tissues (prostate, bladder, lymph node and the seminal vesicle) were compared to the epithelial cells of the prostate. **B** Representative images of immunofluorescent-stained lymph node tissue and bladder for SR cells (cytokeratin, green), cytotoxic T cells (CD8, pink), the programmed death protein 1 (PD-1, yellow) and the nucleus (DAPI). **C** Representative image of the bladder tissue with SR cells (cytokeratin, green) and the programmed death protein ligand 1 (PD-L1, yellow). **D** Proposed model of SRCC DNA damage repair mechanisms and replication stress response.

To confirm our hypotheses derived from our proteomic analyses, which pointed to tumour immunogenicity and DNA damage response pathways, we immunostained for PD-1 and CD8-positive cytotoxic T cells in bladder tissue (primary tumor site), and in lymph nodes (metastasis) (Figure 5B). These tissues exhibited stubstantial or moderate infiltration of PD-1+ cytotoxic T cells, respectively (Figure 5B). We also observed a pronounced upregulation of programmed cell death ligand protein 1 (PD-L1) on SR cells of the bladder (Figure 5C). This suggests that immunotherapy, particularly PD-1/PD-L1 inhibitors, could be a promising therapeutic approach for targeting these tumors.

In line with our findings, the PD-1 inhibitor pembrolizumab had indeed been recommended and administered as a therapy following recurrence rather than chemotherapy. Our results indicate that the ladder, would have been unlikely to be effectivel while having the usual adverse effects. Initiated in 2022, the pembrolizumab ICB therapy on our patient has successfully halted tumor progression, with MRI scans conducted quarterly confirming tumor stasis.

Based on our results we propose a model is which the DNA damage repair mechnisms and the replication stress response takes center stage in SRCCs (Figure 5 D): These SR cells hyper-activate the epidermal growth factor receptor (EGFR) pathway with subsequent hyper-proliferation. Increased DNA replication combined with defective MMR then results in numerous unrepaired post-replication DNA lesions across the genome. The repair of these lesions relies on the cells’ excision repair mechanisms, including base excision repair (BER) and nucleotide excision repair (NER), which we observed to be upregulated at the protein level. The abundance of such lesions, along with the increased rate of DNA replication, are major driving forces behind replication stress, leading to the activation of the ATR signaling pathway. Proteomics indicates SRCC cells respond to this stress by upregulating proteins mediating stalled replication fork protection and collapsed replication forks repair, striving to maintain their genome integrity.

## Discussion

In this study, we employed Deep Visual Proteomics (DVP) to investigate the proteome landscape of Signet Ring Cell Carcinoma (SRCC) across primary and metastatic sites from a single patient. Our analysis of approximately 50 SR cells per organ yielded up to 7,700 proteins, providing unprecedented insights into the tumorigenic properties and potential signaling pathways of SRCC.

We identified both shared and organ-specific protein patterns in SR cells, with a clear distinction from normal epithelial control cells. Key drivers of this difference include mucins, CLCA1, CEACAM5, and CEACAM6. Mucins, particularly MUC1, MUC2, and MUC13, showed significant enrichment in SR cells across all organs, directly contributing to the characteristic signet ring morphology^56^. CLCA1, closely linked to mucin production, can significantly alter the tumor microenvironment, affecting cell adhesion and migration ^36,57^. The upregulation of CEACAM5 and CEACAM6, immunoglobulin-related glycoproteins and adhesion molecules, is notable. While CEACAMs are known to facilitate cellular connection and are frequently elevated in various cancers ^58–60^, their specific role in SRCC has not been previously emphasized. Their overexpression may contribute to the distinctive morphology and aggressive behavior of SRCC through promotion of invasion and metastasis^61–64^.

Our data revealed significant alterations in DNA damage response (DDR) pathways across SR cells in different organs. We observed changes in excision repair mechanisms, including DNA mismatch repair (MMR), base excision repair (BER), and nucleotide excision repair (NER). The upregulation of most MMR proteins, coupled with the downregulation of MLH1, suggests a defective MMR pathway, consistent with previous studies linking microsatellite instability to colorectal SRCC ^17,18^.

A key finding of our study is the upregulation of the ATR signaling axis, indicating ongoing replication stress – a recognized hallmark of cancer driving genome instability^65^. Our proposed model suggests that replication fork stalling and collapse result from an increasing load of post-replicative lesions combined with increased proliferation and DNA replication rates. In response to this stress, the ATR signaling pathway is activated, triggering mediators of stalled replication fork protection, and collapsed replication fork repair and restart (Figure 4G, 4F, 5D)^66^. Single-ended double-strand breaks, the most deleterious form of DNA lesions formed upon replication fork collapse, are addressed by the break-induced replication (BIR) pathway^67,68^. We demonstrated that BIR is upregulated in SR cells across all four organs examined. The error-prone nature of BIR has been associated with high mutation rates, gross chromosomal rearrangements (GCRs), and loss of heterozygosity, further fostering genomic instability^50,69^.

In line with this model, SR cells exhibited a considerable increase in the abundance of poly (ADP-ribose) polymerase (PARP), a key player in DNA repair, and a decreased abundance of APOBEC (apolipoprotein B mRNA editing catalytic polypeptide-like) enzymes compared to adjacent non-tumorigenic epithelial cells. This protein profile suggests a complex interplay between DNA damage accumulation and repair mechanisms in SRCC.

The activation of these DNA damage response and repair pathways likely contributes to the high mutation rate and genomic instability observed in SRCC. This genomic instability, particularly the disruptions in the MMR pathway, is linked to microsatellite instability, which in turn can lead to increased tumor immunogenicity. These findings provide a mechanistic explanation for the observed enrichment of immune-related protein signatures in our proteomics data, and the potential efficacy of immunotherapy in SRCC^17,18^, which we could confirm by immunofluorescence (IF) imaging, revealing strong cytotoxic T lymphocyte infiltration and PD-1 expression^22^. Moreover, we observed alterations related to the complement cascade pathway do occur in SRCC tissues as previously reported^70^

Our results provide a rationale for the observed clinical response to pembrolizumab immunotherapy in this patient^55,71–73^, despite initial sequencing results showing only 0.8% unstable microsatellite sites. This highlights the potential of proteomic analysis in guiding treatment decisions, especially in cases where genomic data alone may not fully capture the tumor’s biology. The identification of replication stress as a central feature of SRCC opens new avenues for targeted therapies. Our findings suggest that targeting the ATR pathway or exploiting vulnerabilities in DNA repair mechanisms could be promising strategies. Additionally, the overexpression of CEACAMs points to potential targets for antibody-drug conjugates or other targeted therapies.

This study demonstrates the power of spatial proteomics in uncovering the molecular intricacies of rare cancers like SRCC. By providing a comprehensive view of the proteome across different organs, we’ve identified common features of SR cells that transcend their tissue of origin, as well as organ-specific adaptations. This approach offers valuable insights into tumor biology that may not be apparent from genomic or transcriptomic analyses alone. In conclusion, our DVP-based analysis of SRCC reveals a complex interplay of DNA damage response, replication stress, and immune signaling pathways. These findings not only deepen our understanding of SRCC biology but also suggest potential therapeutic strategies. The success of pembrolizumab in this case, explained retrospectively by our proteomic data, underscores the potential of precision oncology approaches guided by comprehensive molecular profiling.

Future studies should aim to validate these findings in larger cohorts of SRCC patients and explore the therapeutic potential of targeting the pathways identified here. Moreover, integrating proteomic data with genomic and transcriptomic profiles could provide an even more comprehensive understanding of SRCC biology, potentially leading to improved diagnostic and therapeutic strategies for this aggressive cancer subtype.

## Acknowledgements

The authors thank L. Drici (NNF CPR Proteomics Program) for technical assistance. We acknowledge Richard Denis Maxime De Mets from the Core Facility of Integrated Microscopy for microscopy support. We also thank S. Adams, A. Mund, F.H. Post, L. Niu, JJ. Wang and E. Krismer for engaging and productive discussions.

## CRediT Author contributions

S. Kabatnik, (Conceptualization: Equal; Investigation: Equal; Formal analysis: Lead; Data Curation: Lead; Visualization: Lead; Validation: Equal; Writing – Original Draft Preparation: Lead; Project administration: Equal).

X. Zheng, PhD (Conceptualization: Equal; Investigation: Equal, Visualization: Supporting, Data curation: Supporting; Visualization: Supporting; Validation: Equal; Writing – Original Draft Preparation: Supporting, Writing – Review & Editing: Equal).

G. Pappas, PhD (Conceptualization: Supporting; Writing – Writing & Editing: Equal)

S. Steigerwald (Data curation: Supporting; Original Draft Preparation: Supporting).

M. Padula, PhD (Conceptualization: Equal; Resources: Lead; Supervision: Supporting)

M. Mann, PhD (Conceptualization: Equal; Supervision: Lead; Resources: Lead; Project administration: Equal; Funding acquisition: Lead; Writing – Writing & Editing: Lead).

## Data availability

The proteomics raw data and quantified files were submitted to the ProteomeXchange Consortium through the PRIDE partner repository (https://www.ebi.ac.uk/pride/) with the identifier PXD053079. Image data will be provided upon request by contacting Sonja Kabatnik at.

## Materials and Methods

### Study design and ethical permission

This is a case study. All experiments were performed on a single individual patient who provided us with FFPE blocks from four organs with SRCC presence: bladder, lymph node, prostate, and seminal vesicle. After consultation with the Nepean Blue Mountains Local Health District, they concluded that ‘there is no need for formal application to the Human Research Ethics Committee’ (HREC). The patient provided full consent as a subject of study (HREC study reference: UTS ETH22-7236), including the provision that the proteomic analysis of signet ring adenocarcinoma will be not followed up by any clinical intervention, and there is ‘no risk to privacy or confidentiality’. Thus, the letter and communication with the Nepean Blue Mountains Local Health District acts as ‘evidence of waiver of the need for HREC approval’.

### Immunohistochemistry and high-resolution microscopy

A detailed protocol for FFPE tissue mounting and staining on membrane PEN slides 1.0 (Zeiss, 415190-9041-000) is provided in the original Deep Visual Proteomics (DVP) article^26^. The tissue sections were initially subjected to deparaffinization and hydration through three cycles involving xylene and decreasing ethanol concentrations from 99.6% to 70%. For Wheat Germ Agglutinin (WGA) labeling, sections on membrane PEN slides were incubated with WGA staining solution (Biotium, 29023; diluted 1:1000) in a light-protected environment at 37°C for 10 min. For pan-cytokeratin (CK), CD8, PD1, PDL1 and pATR staining, antigen retrieval was achieved by immersing the tissue sections on glass slides in EDTA buffer (Sigma, E1161; pH 8.5) at 90°C for 30 min. Following this, the tissue sections were blocked with TBS protein-free blocking buffer (LI-COR, 927-80000) for 20 min at room temperature. For CD8/PD1/CK triple staining, the sections underwent overnight incubation at 4°C with anti-CD8 antibody (Abcam, ab17147; 1:100), followed by slide washing and subsequent incubation with Alexa Fluor® 647 goat anti-mouse antibody (Invitrogen, A-21235; 1:1000) for one hour at room temperature. After rinsing, the slides were further incubated overnight at 4°C with anti-PD1 antibody (Miltenyi Biotec, 130-117-384; 1:100) and anti-CK antibody (Invitrogen, 53-9003-82; 1:500). For PDL1/CK double staining, slides were incubated with anti-PDL1 antibody (Invitrogen, 12-5983-42; 1:100) and anti-CK antibody (Invitrogen, 53-9003-82; 1:500) overnight at 4°C. For pATR staining, slides were incubated with anti-pATR antibody (GeneTex, GTX128145; 1:500) overnight at 4°C, followed by slide washing and subsequent incubation with Alexa Fluor® 647 donkey anti-rabbit antibody (Invitrogen, A-31573; 1:1000) for one hour at room temperature. Finally, we used DAPI (Abcam, ab228529; 1:1000) for nuclear counterstaining for 5 min at room temperature and the slides were mounted with Anti-Fade Fluorescence Mounting Medium (Abcam, ab104135) before examination under an AxioScan7 microscope (Zeiss, for WGA, CK, CD8, PD1 and PDL1 imaging) or PANNORAMIC 250 Flash III (3Dhistech, for pATR imaging).

Tumor regions were identified using CK staining, WGA staining, or simply by including the auto-fluorescent signal of mucin.

### Cell segmentation and classification

Microscopy images were imported into BIAS (Biology Image Analysis Software, <u>single-cell-</u> <u>technologies.com</u>), for machine learning-based cell segmentation, classification, and subsequent single-shape export for semi-automated laser microdissection. For SR cell segmentation, we utilized a pre-trained deep neuronal network on our IF WGA-stained tissues. Detection confidence was set to 60% and the contour confidence to 20%. Cell shapes with a larger area than 1000 µm^2^ were excluded. To accurately classify SR cells, we trained a BIAS-integrated multilayer perceptron (MLP) feedforward neural network on manually identified SR cells across all four tissues. We set the weight scale and the momentum parameter to 0.01, and the number of iterations to 10,000. Subsequently, reference points were set, and SR cell contours were exported for semi-automated laser microdissection^26^.

### Laser microdissection

After aligning the reference points using the LMD7 (Leica) microscope, we imported the shape contours to facilitate semi-automated laser microdissection, which was conducted with the following parameters: laser power at 34, aperture set to one, cutting speed at 28, the middle pulse count to tree, final pulse to one, head current at 47 percent, pulse frequency at 2,600 Hz, and an offset of 180. For each type of organ tissue, SR cell shapes were excised in triplicates, and collected into 384-well plates, deliberately omitting the outermost rows and columns. After microdissection, we spun down the plate at 1,000 g for 10 min, and the dissected cell shapes were preserved by freezing at –20°C for later processing.

### MS sample preparation

The entire MS sample preparation protocol was adapted from the original DVP paper ^26^. After protein digestion, samples were vacuum dried, resuspended in 20 µL Evosep buffer A (0.1% formic acid v/v) and directly loaded on Evotips (https://www.evosep.com/).

### LC-MS

Subsequently after Evotip loading, our low input samples were analyzed on our Orbitrap Astral mass spectrometer (Thermo Fisher Scientific) connected to the EvoSep One chromatography system (https://www.evosep.com/). We utilized a commercial analytical column (Aurora Elite TS, IonOpticks) and an EASY-Spray™ source to run our samples with the 40 Samples Per Day (’40 SPD’) method (31-min gradient). All samples we recorded in DIA (data independent acquisition) mode. The Orbitrap analyzer of the mass spectrometer was utilized for full MS1 analyses with a resolution setting of 240,000 within a full scan range of 380 – 980 m/z. The automatic gain control (AGC) for the full MS1 was adjusted to 500%. For the acquisition of our low-input FFPE DVP samples, we set the MS/MS scan isolation window to 3 Th (200 windows), the ion injection time (IIT) to 5 ms, and the MS/MS scanning range to cover 150 − 2000 m/z. Selected ions were fragmented by higher-energy collisional dissociation (HCD)^74^. at a normalized collision energy (NCE) of 25%.

### MS data analysis

Raw files were first converted to the mzML file format using the MSConvert software (https://proteowizard.sourceforge.io/) from Proteowizard, keeping the default parameters and selecting ‘Peak Picking’ as filter. Afterwards, mzML files were quantified in DIA-NN^28^ (version 1.8.1) using the FASTA (2023, UP000005640_9606, with 20,594 gene entries) from the UniProt database and a direct-DIA approach. The enzyme specificity was set to ‘Trypsin/P’ with a maximum of two missed cleavages. Parameters for post-translational modifications were set to including N-terminal methionine excision, methionine oxidation and N-terminal acetylation were all activated, and a maximum of two variable modifications were allowed. Precursor FDR was set to 1%, and both mass and MS1 accuracy were set to 15 ppm. ‘Use isotopologues’, ‘heuristic protein inference’, ‘no shared spectra’ and ‘match between runs’ (MBR) were enabled. Protein inference was set to ‘genes’ and the neural network classifier run in ‘single-pass mode’. We chose the ‘robust LC (high precision)’ as quantification method and a retention time-dependent cross-run normalization strategy. SRCC samples, microdissected across all four organs’, were searched together.

### Bioinformatic analysis

After quantification in DIA-NN, the protein group matrix was imported into Perseus^75^, and samples were annotated according to the organ of origin (bladder, lymph node, prostate, and seminal vesicle). Proteins with 70% of quantitative values present ’in at least one group’ were then kept for imputation of missing values based on their normal distribution (width = 0.3; downshift = 1.5). Further, all statistical tests were corrected for multiple hypothesis testing, applying a permutation-based false discovery rate (FDR) cut off either 5% or 1%.

Gene Set Enrichment Analysis (GSEA) was conducted using Python (version 3.9.7) and the GSEApy package (documentation: https://github.com/zqfang/GSEApy, version 1.0.4).

For the purpose of data visualization, our analyses were performed using the Python programming language (version 3.9.7), and essential libraries such as NumPy (version 1.20.3), Pandas (version 1.3.4), Matplotlib (version 3.4.3), and Seaborn (version 0.12.2). Additionally, the ShinyGo web tool (documentation: http://bioinformatics.sdstate.edu/go/), version 0.77, was used to perform gene ontology (GO) term enrichment analysis.

